# Genetic architecture and adaptation of Ladakh highlanders of trans-Himalayas

**DOI:** 10.1101/2024.02.05.579041

**Authors:** Lomous Kumar, Richa Rajpal, Bhavna Ahlawat, Nagarjuna Pasupuleti, Snigdha Konar, Aparna Dwivedi, Sachin Kumar, Sonam Spalzin, Stanzen Rabyang, Kumarasamy Thangaraj, Niraj Rai

## Abstract

Trans-Himalayan Ladakh has witnessed complex cultural movements and demographic changes since the Neolithic period, which is still continue despite the harsh, inhospitable and cold climate. Although geographically isolated from mainland South Asia, Ladakh has historic trade routes and is well connected and accessible to travelers from Tibet and Central Asia. Despite its rich cultural heritage, a detailed description of the genetic landscape of the Ladakh region is completely lacking, particularly with regard to genome-wide analysis and larger sample sizes. Therefore, in the current study, we genotyped 80 individuals from Kargil and Leh districts of the Union Territory of Ladakh, India. Here, we performed a comprehensive genetic analysis based on allele frequency and haplotype sharing. Our analysis revealed the presence of two distinct genetic lineages in the region with quite distinct genetic composition. The population of Leh Region is more similar to East Asian and Southeast Asian populations. In contrast, the population of the Kargil Region (LDKLA) is more similar to Indo-European populations. Demographic modeling suggests that the Leh group shares a genetic history with Tibetans, while the Kargil group showed great affinities with Kashmiri Muslims, Gujjars and Nepalese Brahmins, and both showed recent admixture. Both groups have experienced a founder event around during 11th to 22nd generations ago, the duration of which coincided with the Mughal invasion. The genome-wide scan for a signal of positive selection revealed genetic signatures of high-altitude adaptation (*EPAS1* and *ELMO2*) in the Leh population, while in the Kargil population the key gene signatures were associated with immunity and female fertility.

## Introduction

Ladakh, the India’s ice-cold desert, is an isolated area in the Indian outback often referred to as Little Tibet, the land of high passes. The passes in this region allowed people to establish trade routes and strategically connected the region with many routes. Despite the isolation, the passes in Ladakh allowed people to establish trade routes from Tibet and Central Asia. This made the region strategically important for important trade routes such as the Silk Road. It is the intersection of several roads passing through Hoshiyarpur, Kullu, Rohtang Pass, Keylong, BaraLacha Pass, Lahchung Pass, Tanglang Pass, Srinagar, and Karakoram Pass to reach Amritsar from Yarkand ^1^. The ancient trade routes played a crucial role in the flow of ideas, people, cultures, exotic goods, and religions. Along these routes, monasteries established positions as mentors, promoted trade, and sent funds to missionaries and traders ^1–3^. Archaeological expeditions explored Ladakh’s past, focusing on ancient monasteries, abandoned trade routes, and enigmatic petroglyphs. Ladakh, known as the Roof of the World, was the site of the first prehistoric settlement and was inhabited by diverse people ^4,5^. The discovery of Paleolithic and Neolithic artifacts in this region indicates that early humans once lived there ^5,6^. The Paleolithic artifacts showed at lEast three phases of cultural development, with the Lower Paleolithic representing the initial phase and the Middle Paleolithic representing the final two phases ^7^. Research along the banks of the Indus River has revealed several Paleolithic tools, including discoids, worked stones, one- and two-sided choppers, and primary flakes ^5^. The Neolithic sites of Burzahom and Wakha have remains of two different types of Kashmir’s earliest pottery ^8^. There are also two different types of pottery in Ladakh: the first type is found in Changthang, Upper Ladakh and Zanskar (characterized by cord-shaped imprints) and the second type is found in Leh (painted pottery more similar to Western Tibet) ^8^.

There is a large gap between the Neolithic period and the Kushan Empire in the first century AD. Before the spread of Buddhism, Ladakh is said to have been dominated by the Bon religion, which existed until the 7th century BC ^5^. It is generally accepted that Sthavira Madhyantika and other fellow monks wrote the preface to Buddhism in Ladakh ^5,9^. Therefore, there are numerous archaeological sites in Kargil, which forms an important Buddhist center in Ladakh. However, there still remains a gap in the genetic and archaeological evidence of human occupation of Ladakh. Genetics show the influx of important maternal ancestors from China into the Tibetan Plateau during the Upper Paleolithic and early Neolithic ^10^. Mitochondrial DNA markers have confirmed that the majority of maternal lineages with additional archaic Denisovan-like autosomal elements arrived on the Tibetan Plateau from Northern China during the latest Upper Paleolithic and early Neolithic (Middle Holocene) ^11,12^. The historical evidence for the occupation of the area by the Kushan Empire in the first century CE, in the mid-7th to mid-9th centuries CE by the Tibetans, and in the 14th to 16th centuries CE due to frequent Mughal invasions are the responsible factors for the increase in genetic mixing of Ladakh populations ^13,14^.

Ladakh, along with Kashmir, was conquered by Muslim invaders in the 14th century, which was later followed by repeated invasions called jihad ^15^. The decentralization of the Mughal Empire, relocated from the mountains of Ladakh and the Hindu Kush, formerly called *Upariyena* (in Vedic Sanskrit) or *Upirisana* (in Avestan), comes from Proto-Iran and means covered with juniper, so out of reach of eagles ^16^. The naming of the mountain as Hindu Kush came after the genocide of Hindu slaves during transport to Central Asia between the 9th and 18th centuries. This all led to gene flow from South Asia to Ladakh and the Trans-Himalayas ^17^. The local demography of the Ladakh region reflects a co-occurrence of people with distinct customs and cultures. The Dah-Hanu area of Leh is inhabited by a unique Dard-speaking tribe that claims its origins in the army of Alexander ^18^. These tribes are called Brokpa, Brogpa, or Broqpa. The term Brok means hill, and pa is the people; so Brokpa are the mountain people, the highlanders. They mainly follow Buddhism and have the traditional *Phasphun* or clan system ^19,20^. The cremation culture and speaking of the Dardic language, Brokskat, suggest that they are descendants of the protorigvedic culture ^21^. The tribe became popular because there were rumors that they were the last pure Aryans ^22^. The pastoral, semi-nomadic Changpa tribe was one of the region’s main inhabitants throughout the Neolithic period, which was also marked by community migrations. The tribe is believed to have originated on the Tibetan Plateau ^7,23^. Changpa or Champa inhabit the cold deserts between Ladakh and Tibet, which are extremely cold and plagued by snowstorms. They converse in Changskhat, a Tibetan dialect. The Changpa people are sometimes referred to as the Changthang Plateau people, who settled the area around the 8th century and came from Tibet. They mainly follow Buddhism ^24,25^. Another tribe, the Monpa people, is believed to have originated from the Monyul region, which existed South of Tibet between 500 and 600 AD ^26^.

Genetic studies of trans-Himalayan migration have resulted in conflicting views, with some viewing the Himalayas as a natural barrier to human migration while others view them as beneficial ^27^. Ladakh populations have been shown to have a close relationship with Southeast Asia ^28^, highlighting that the populations from Ladakh were not genetically isolated. The Y chromosome diversity in Ladakh is very high and has a mixed prevalence of East, West, and South Asian Y haplogroups. This also confirms the fact that the region lies at a genetic confluence as it has experienced many episodes of demographic change with cultural and genetic changes due to trade, immigration, migration, and military interventions ^28,29^. The abundance and diversity of mitochondrial haplogroup M9 highlight the potential genetic contribution of the Tibetan Plateau and China ^30^. This genetic proximity to East Asians is also supported by the frequency of Hg A, a haplogroup not so common in India but very common among Tibeto-Burmans, including Tibetans and Mongolians ^28–32^. Genetic studies carried out to date are based on a limited number of samples and low-resolution markers. The present study is the first attempt to gain deeper insight into the origin and genetic relationship of Ladakhi genomes. We performed an in-depth autosomal analysis of the Brokpa, Changpa and Minaro populations.

## Methodology

### Sampling

A total of 80 unrelated Ladakh individuals were enrolled in this study, which comprised Minero (n = 11), Brokpa (n = 35) from Kargil district, and Changpa (n=34) from Leh district in the Union territory of Ladakh. All subjects confirmed in a questionnaire that they and their parents were born in Ladakh and had lived there their entire lives. Genomic DNA was extracted from venous blood using the standard phenol-chloroform method. This research was approved by the Ethics Committee of the Birbal Sahni Institute of Palaeosciences, Lucknow. All experiments were carried out in accordance with relevant guidelines and regulations. All participants provided written informed consent before beginning this study.

### Genotyping and Quality control

Genotyping of all DNA samples was performed using Illumina HumanOmniExpress-24-v1 at a total of 642,824 genome-wide single nucleotide polymorphism (SNP) sites. Changpa samples came from four different locations and were referred to as LDK_BS, LDK_KDL, LDK_VA, and LDK_V, Minero as LDK2_RK; and Brokpa as LDK_RK genotype samples in the final dataset. All genotyped Ladakh individuals were merged with modern references from the Simon Genome Diversity Panel (SGDP) ^33^, the Human Genome Diversity Panel (HGDP) ^34^, 1000 Genomes Panel ^35^ and, unpublished South Asian genotype data from our laboratory. The combined data was quality filtered using Plink-1.9 ^36^ and with a quality cutoff of geno 0.03 and mind 0.05. For principal component analysis and admixture, the combined genotype data were adjusted for linkage disequilibrium (LD) (independent pairwise 200 25 0.4). For haplotype-based analysis and Identity-By-Descent (IBD) analysis, reference groups with fewer than five individuals were excluded in the downstream analysis. SGDP samples (due to low sample size per population) were excluded in the haplotype-based analysis and all F-statistic calculations to increase the power of the statistics by increasing sample numbers.

### Clustering analysis

The clustering analysis for combined genotype data was performed using Principal Component Analysis (PCA) and Admixture analysis. PCA was done using *smartpca* tool in EIGENSOFT^37^ package. The first two principal components were plotted using custom R script. Admixture analysis was done using a Admixture v1.3.0 ^38^ by first calculating the cross-validation (CV) error from K = 2 to K = 12. Then final Admixture run was performed at the K value with the lowest cross-validation error, and Q-matrix was plotted using a custom R script.

### Three population and four population tests and Maximum Likelihood based tree

Three population tests were performed using Admixtools 2 ^39^ to calculate shared genetic drift in the form of outgroup F3 statistics and Mbuti as the outgroup population. Gene flow pattern was inferred using the qpDstat function of Admixtools 2 ^39^ package in R. All the results for outgroup F3 tests were plotted using custom R scripts. A Maximum-Likelihood tree was constructed in TreeMix v-1.1 ^40^ using the outgroup population as Mbuti and with a LD window of 500 SNPs. Tree was plotted using plot_tree.r script provided with the package.

### Haplotype based clustering and Identity by Descent analysis

We also used ChromoPainter-v2 ^41^ and FineStructure-v4 ^41^ to infer fine-level population clustering and recent gene flow by haplotype sharing pattern using a haplotype-based approach. We first phased our data with SHAPEIT5 ^42^ using default parameters. Chromosome painting was performed using ChromoPainter^41^, first by doing 10 EM iterations with 5 randomly selected chromosomes with a subset of surrogate and target individuals to infer the global mutation rate (µ) and switch rate parameters (Ne). Then, we performed the main ChromoPainter run for downstream FineStructure ^41^ analysis with a switch to paint all recipients with all donor haplotypes using an estimated fixed value of the global mutation rate and switch rate parameters. FineStructure was run using an inferred chunk sharing matrix and 5 million iterations with 1 million burns in each iteration. The final tree was plotted using the provided R script in the FineStructure-v4 package, with some modifications.

For identity by descent analysis, we performed haplotype inference or phasing with three independent runs of Beagle-5.4 ^43^. IBD segments were determined from phased data of all three runs separately using refined-IBD ^44^, subsequently combining segments from all three runs, and then combined segments were merged using the merge-IBD-segments tool. Then the IBD sharing matrix was plotted using a custom script in R. Principal component analysis was performed using the Chromopainter matrix, and the first two components were plotted using a custom R script. Runs of homozygosity in two window sizes (1000 kb and 5000 kb) were calculated using Plink-1.9 ^36^.

### Genetic simulation to test the effect of founder event on RoH distribution

We performed genetic simulations using msprime-1.3.1 ^45^ to reconstruct the founder event scenario of both the Ladakh clusters. For all simulations, we started with a population of size 12500 at 1800 generations (similar to the African split) and produced 60 sample VCFs. We used a genome size of 50 Mb with a uniform recombination rate of 1e-8 and a mutation rate of 2.36e-8. For the LDK-LA-BOT population scenario we used the founder event strength similar to LDKLA (Ladakh Low Altitude; I_f_ = 2.1, Tf = 11-13 gen) group. For LDK-HA-BOT population scenario, we used the founder event strength similar to the LDKHA (Ladakh High Altitude; I_f_ = 1.1, Tf = 21-24 gen) group. Founder event in simulated samples were tested using ASCEND-10.1.1 ^46^. Runs of homozygosity were performed on simulated data (VCF format) in Plink-1.9 ^36^ and plotted in R using a custom script.

### Admixture modelling with modern Eurasians and Demographic model inference

In order to infer best fitted admixture model and time of admixture using modern Eurasian source populations, we used haplotype-based method implemented in fastGlobeTrotter ^47^. We first performed phasing using SHAPEIT5 ^42^, then performed chromosome painting using Chromopainter2 ^41^. This was done in two steps: in first step, copy vectors files were generated using all donor and recipient haplotypes, and in second step, sample files for target haplotypes were produced using putative sources of admixture as surrogates. Finally using combined chunk length file and sample file fastGlobeTrotter run was performed using 10 mixing iterations for each Ladakh groups (LDKHA and LDKLA) separately.

In order to infer demographic history of LDKHA and LDKLA groups, we used Moments’ ^48^ inference optimize function in python. We constructed five alternate models for both LDKHA and LDKLA groups using Demes ^49^ python package, describing the admixture scenario of fastGlobeTrotter. We selected the best fitted model for both Ladakh groups based on best Loglikelihood inference. Best fitted demographic parameters were inferred for this model using Moments and confidence intervals were calculated using *moments.Demes.Inference.uncerts* function.

### Cross population test for genome wide signatures of selection

We performed a cross-population test for selection in (i) the high-altitude (11,480 ft) Ladakh population (LDK_BS, LDK_KDL, LDK_VA, and LDK_V), jointly referred as LDKHA and (ii) the lower-altitude (8,780 ft) Ladakh group (LDK_RK and LDK2_RK), jointly referred to as LDKLA, using the R package *rehh* ^50^. We phased the genotype data using SHAPEIT5 ^42^ and then used the phased haplotypes of LDKHA and LDKLA with a minor allele frequency cutoff of 0.05 to perform an XP-EHH selection test with the East Asian and European population, respectively, for the reference population. These reference groups were selected based on the genetic affinity inferred from PCA, admixture, and haplotype sharing. The candidate regions were determined using a threshold value of 8, and a window of 100kb was used with 10kb overlaps. The flanking region genes were determined using a custom R script and a human genome annotation file (GFF3 format). Manhattan plots for genome wide regions depicting selection signatures were done with custom R scripts.

## Results

### Genetic structure of Ladakh highlanders

In the allele frequency-based principal component analysis, all Eurasians were distributed along three major axes, with one extreme formed by West Eurasian groups like Caucasus and European populations, another extreme occupied by East Eurasian populations from East Asia and South East Asia, and in the third axis, South Asian Austroasiatic and Dravidian linguistic groups were present (Fig. 1b). Ladakh populations were split into two genetically distinct clusters. Four Ladakh groups from the Leh region (LDK_BS, LDK_KDL, LDK_VA, and LDK_V) were forming an extended genetic cluster along the East Asian and South-East Asian axis, but not overlapping with either East Asian or South-East Asian groups. The second cluster of two Ladakh groups (LDK_LK and LDK2_RK) was more into the Indian main cline and more specifically among the North (Jatav, Bihar, and Jain) and North-West Indian (Kashmiri-Muslim, Khatri, and Gujjar) Indo-European groups and also Nepal-Brahmins (Fig. 1b). The first group closer to East and South-East Asians is mainly comprised of the Changpa population from the Leh region, while the second group closer to Indo-Europeans is comprised of the Minero and Brokpa populations from Kargil region. Few individuals from Brokpa and Minero, as well as Nepal Brahmins, lie in between two Ladakh clusters.

**Fig 1.**
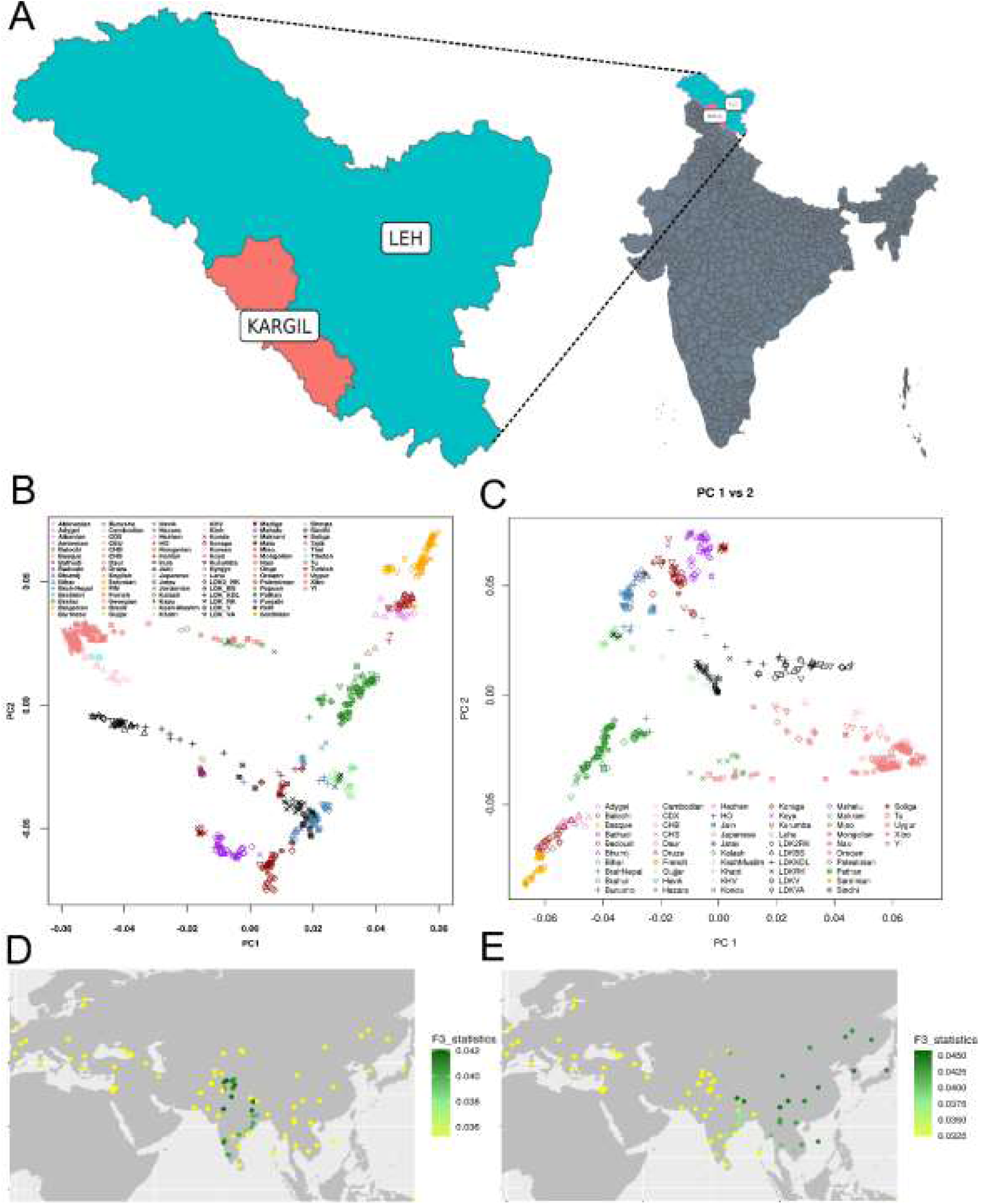
**A.** Sampling locations of Leh and Kargil districts in the union territory of Ladakh. **B.** PCA biplot with allele frequency-based analysis, **C.** PCA biplot with Chromopainter Chunk sharing matrix, **D.** Outgroup F3 statistics of Brokpa and Minero with all Eurasians, D. Outgroup F3 statistics of Changpa with all Eurasians. (**LDKLA = LDKRK and LDK2RK; LDKHA = LDKBS, LDKKDL, LDKVA and LDKV**)

In our model-based clustering using nine ancestral sources (K = 9), we observed a very clear and contrasted distribution of ancestral components between two Ladakh groups, referred to as LDKHA (Changpa) and LDKLA (Brokpa and Minero) hereafter. LDKHA group is maximized in blue color component, which is also present majorly among Tibetan, Hazara, Uyghur, East and Southeast Asians and with minor presence in LDKLA, Nepal Brahmin and Burusho populations (Fig 2a). LDKLA group is maximized in the forest green-colored ancestral component, which is present in Nepal Brahmin, Kashmiri Muslim, Burusho and Khatri populations but minor presence in LDKHA and Tibetan. So, both the Ladakh groups have minor presence of each others’ major ancestral component. However, Kashmiri Muslim only share component with LDKLA but not with LDKHA. A high amount of West-Eurasian component was observed in four Brokpa (termed LDKLAw; w for west Eurasian) individuals (typical of North-West Indian populations), consistent with PCA-based clustering. On the contrary some individuals of LDKLA group shows higher presence of blue component typical of LDKHA, hence termed LDKLAE (E for East Eurasian) (Fig 2a).

**Fig 2.**
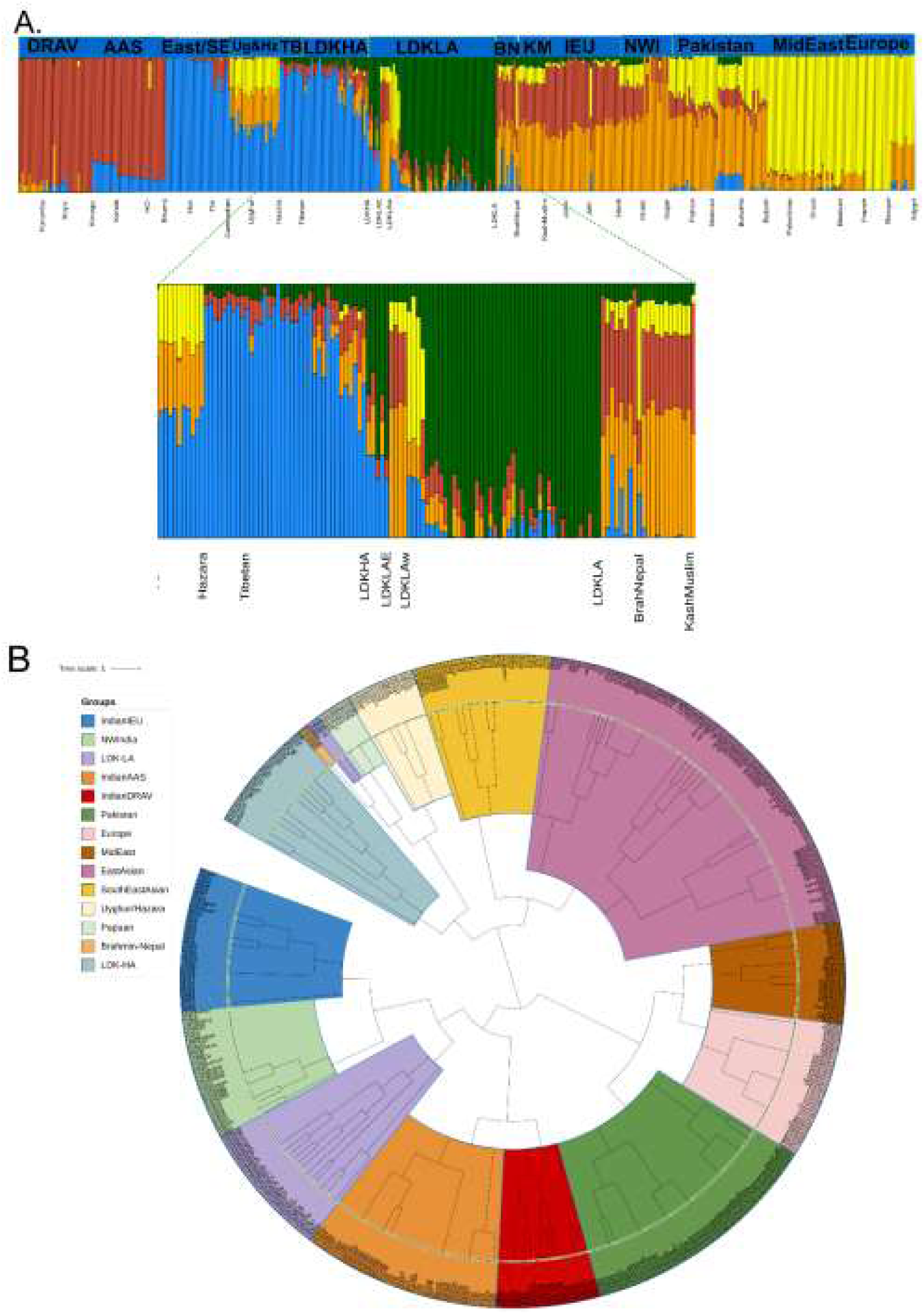
**Admixture bar plot** of Ladakh populations with modern Eurasians and expanded regions highlighting affinity of Both Ladakh clusters with geographically proximal populations **A. FineStructure ML tree** showing haplotype-based clustering of both Ladakh clusters (highlighted with separate colors **B. (Group legends are anticlockwise)**

### Haplotype based genetic structure

In the haplotype sharing based genetic structure inferred in the form of the FineStructure ^41^ tree, all Eurasian populations were distributed between two major clades, one occupied by West Eurasian populations and the other by East Eurasian populations (Fig 2b). Consistent with the frequency-based clustering, the LDKHA group was sharing the clade with East Asian and South-East Asian populations, while LDKLA was present in the West Eurasian clade along with Indian Indo-European populations, Pakistan, the Middle East, and European populations (Fig. 2b). However, four LDKLA and three Nepal Brahmins were present in the East Eurasian clade, along with Hazara, Uyghur, and Papuan populations (Fig. 2b). Compared to frequency-based PCA, Chromopainter haplotype-sharing matrix-based PCA revealed a very tight clustering pattern of the LDK_RK (Brokpa) and LDK2_RK (Minero) groups with the Gujjar population (Fig. 1c). Whereas, four Brokpa individuals were still clustered with Kashmiri Muslim individuals in the main Indian cline, indicating their very close genetic affinity. On the other hand, Changpa (LDK_BS, LDK_KDL, LDK_VA, and LDK_V) were making distinct clusters with few individuals in between both Ladakh clusters, indicating some shared genetic history (Fig. 1c).

### Gene flow pattern inferred from F-statistics

In the outgroup F3 statistics, a distinctive pattern of allele sharing was clearly observed. LDKLA (Brokpa and Minero) groups had the highest outgroup F3 statistics distribution (represented by a darker green color), with mostly North-West Indian and North-Indian Indo-European populations (Fig. 1d). LDKHA (Changpa) shared the highest drift with Tibetan, Sherpa, East Asian, and South-East Asian populations (Fig. 1e). In Patterson’s D-statistics ^51^ in the form of F4 (Mbuti, LDKHA/LDKLA: East Asian/South East Asians, West Eurasian/Indian Indo-Europeans), to infer gene flow pattern in Ladakh population, we observed a similar pattern with both Ladakh groups as observed in outgroup F3 statistics. LDKHA groups had higher gene flow from East Asian, South East Asian, Tibetan, and Sherpa (negative D-statistics) populations compared to most of the West Eurasian and South Asian groups (Supplementary Table S1). Whereas, LDKLA groups showed higher gene flow from Indian Indo-Europeans and West Eurasian populations (positive D-statistics) except Sindhi, Makrani, and Balochi from Pakistan (D-statistics were negative with these groups) (Supplementary Table S1). We considered only those D-statistics which that had a z-score higher than 3 in all cases.

### Identity By Descent segments sharing and Runs of Homozygosity (RoH)

The genome-wide distribution patterns of Runs of Homozygosity segments in the 1 kb window size showed a higher distribution in LDKLA groups in terms of mean number of segments compared to LDKHA groups (Fig. S3a), whereas, in terms of mean total length both the groups were almost comparable. In the higher window size of 5 kb, three of the LDKLA groups (LDK_RK and LDK2_RK) are outcompeting the LDKHA (LDK_BS, LDK_KDL, LDK_VA, and LDK_V) groups (Fig. S3b).

In terms of intra-population IBD sharing, Brokpa (LDK_RK) and Minero (LDK2_RK) had very high IBD segments sharing within the population (with an LOD score > 10) and were comparable to the Gujjar population (Fig. 3a). All the Changpa groups (LDK_BS, LDK_KDL, LDK_VA, and LDK_V) showed comparatively lower intrapopulation IBD segments sharing (with an LOD score > 10). In the cross-population IBD sharing, both Ladakh groups share the highest number of IBD segments with each other than with other Eurasian populations (Fig 3b). Ladakh-LA group (Brokpa and Minero) share more IBD segments with Kashmiri Muslims, while Ladakh-HA (Changpa) shares more segments with East Asian populations (Yi and Tu) from HGDP panel (Fig 3b).

**Fig 3.**
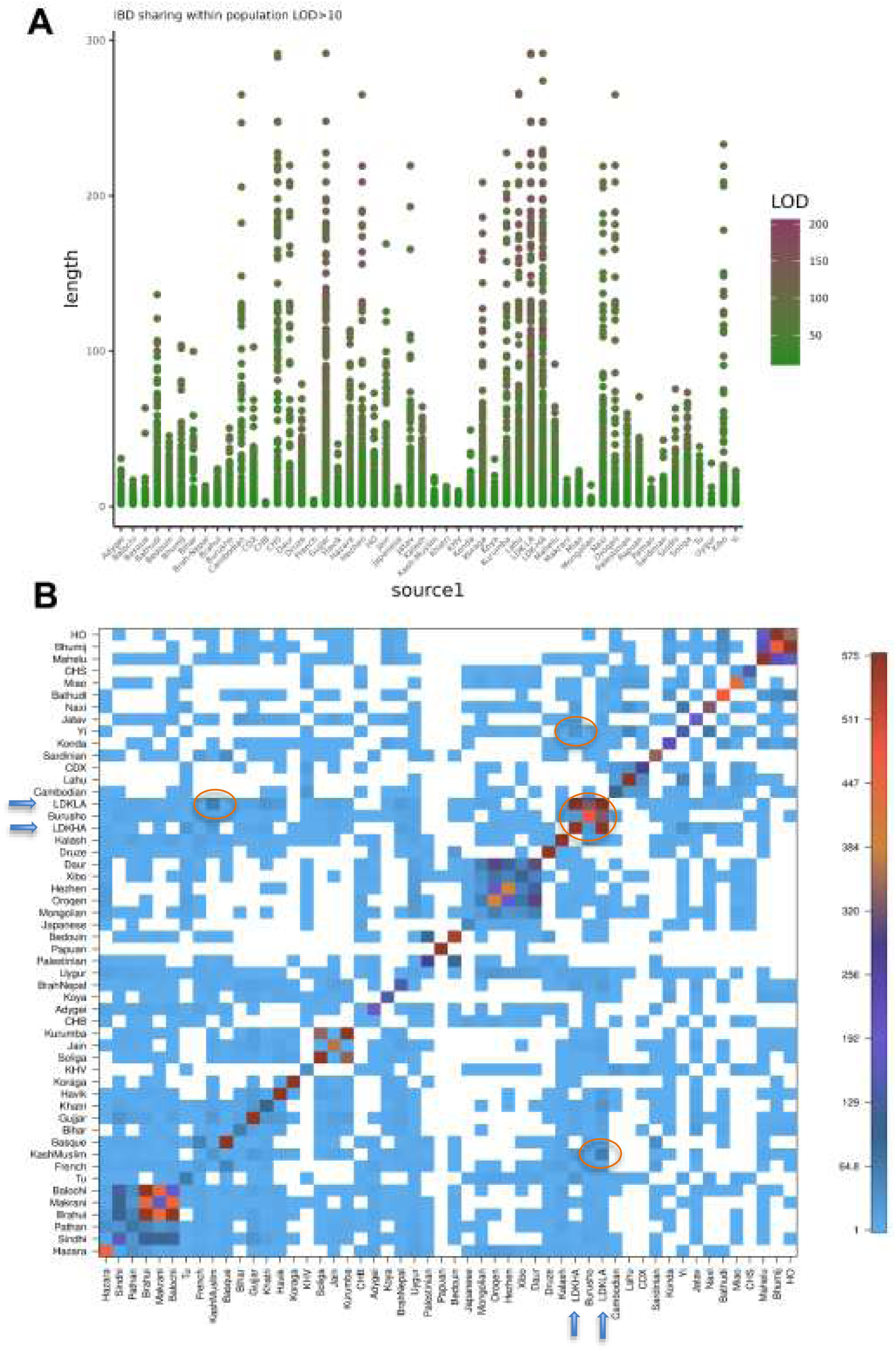
Intra-population **A.** and Cross-population **B.** IBD sharing matrix adjusted for sample number. Elipse in lower panel show pattern of sharing)

### Effect of founder events on RoH distribution based on simulated data under founder event scenarios of Ladakh groups

We tested the effect of historical changes in population size on the RoH distribution in the simulated genomes under two different scenarios: (i) a founder event 13 generations ago with a strength similar to that of Ladakh Low Altitude (LDK-LA-BOT), and (iii) a bottleneck 24 generations ago with a similar strength of the founder effect as observed in LDKHA (LDK-HA-BOT). We 50Mb genome size and 20 chromosomes for each genetic scenario and found that the RoH distribution (mean KB and mean NSEG) is similar to the actual observed data (almost comparable distribution in LDK-LA and LDK-HA at 1kb and higher distribution at 5kb in LDK-LA).

### Demographic model and parameter inference

In the fastGlobeTrotter admixture modelling, best fit admixture model for LDKHA (Ladakh High Altitude) group was one date model with surrogates as Tibetan and Kashmiri Muslims (Table S2). The admixture date was 23 generations ago with 80% admixture from Tibetan and 20% from Kashmiri Muslim (Indo-European putative source) (Table S2). For LDKLA (Ladakh Low Altitude) group from Kargil has one date admixture at 27 generations before with surrogates as Kashmiri Muslim (77% admixture proportion) and Tibetan (∼13% admixture proportion).

Based on these admixture models we further explored the demographic history of both LDKHA and LDKLA groups using Moments. We explored five distinct models describing different ancestral source group and later time admixture pulse scenario for both the groups. The best fitted putative ancestral source group was Tibetan for LDKHA group with a recent pulse of admixture from French (proxy for Indo-European like admixture source) (Supplementary Fig 1a-b). The Log-likelihood value for this model was - 114400.79498136348 with migration rate from Tibetan to LDKHA equal to 0.0379. This model shows significant decrease in Tibetan effective population size (from 9.99e+04 to 1.36e+03), while slight increase in case of LDKHA (from 100 to 462).

The best fitted putative source group in case of LDKLA was derived from an admixture between Tibetan and French like group (approximately 100 generations before present) (Log-likelihood - 121125.18700491334) and an additional pulse of admixture recently from Tibetan (representing admixture of Kashmiri Muslim and Tibetan in fastGlobeTrotter run) source (Supplementary Fig 2a-b). There is slight decrease in effective population size in LDKLA (from 698 to 439) with migration rate of 0.00255 from French like group. The pulse time of admixture from Tibetan was 40 generations before present (95% CI 36.032-44.8412) (Supplementary Fig 1a-b).

### Signatures of genetic selection in Ladakh groups

We performed a cross-population test for inferring genetic signatures of selection in both Ladakh groups (LDKHA and LDKLA) individually. For LDKHA, we used East Asian populations as the reference group, and using the XP-EHH approach, we found a total of 33 Single Nucleotide Polymorphism (SNP) markers under positive selection (Supplementary Table S2, Fig S5a). These markers defined three putative candidate regions on chromosome 2 (16 SNPs), chromosome 3 (5 SNPs), and chromosome 7 (12 SNPs) under selection (Supplementary Table S3, Fig S5a). In total, 22 Gene signatures were found in 250 kb flanking region of three putative candidate regions in the LDKHA group. Some of these are well-known signatures of high-altitude adaptation (*EPAS1, ELMO2*), while many of these genes are broadly categorized under Immunity related genes (Supplementary Table S3). For LDKLA, we used the European population as a reference group in cross population test, and we found a total of 57 genetic signature loci under selection in three candidate regions (two on chr4 and one on chr9) (Supplementary Table S4, Fig S5b). Some of the gene signatures in the flanking region of candidate regions on chr 4 are strongly expressed in blood vessels (*OCIAD1 and 2*), while others are related to immunity and female fertility (*TXK* and *TEC* are immunity-related and *ZAR1* for female fertility) (Supplementary Table S5, Fig S5b).

## Discussion

Ladakh is a cold, high-altitude desert region where cultivation is not profitable without irrigation due to the short growing season and low annual rainfall. The presence of Neolithic sites and historical evidence suggest that suitable climatic conditions for human habitation existed near the Middle Holocene ^7^. Ladakh was one of the most important regions in NorthWest India and was known for Buddhist sacred artifacts, gold, silver, borax, salt, copper, and precious fabrics. The region’s high ingenuity made it one of the recognized main trade routes. These hypotheses are supported by the presence of archaeological evidence that the most important stations along the above-mentioned routes were marked by petroglyphs and massive rock sculptures of Buddhist deities ^1^. The earliest known petroglyphs in the region were created by steppe populations associated with the Chinese Empire and Persia, dating back to the Neolithic or Bronze Age ^52^.

However, despite the archaeological evidence, the exact origin and genetic ancestry of the Ladakhi peoples are still unknown. The exact sequence of migration movements and the timing of their arrival are still a question mark for the scientific community. The land was inhabited by several groups. Previous research in this area has mentioned numerous references to human migrations. Hunters are said to have found the site, planted wood and grain to test the fertility of the land, and eventually built a hamlet or fort ^53^. The Dards of Gilgit and neighbouring Baltistan, also known as the first known people of Ladakh, established it as a Goldilocks zone; however, they were driven out by the Tibetans. It has also been documented that nomadic pastoralist communities from neighboring areas such as Changthang and Zhang-zhung moved to the region seasonally in search of shelter and resources; the group of remaining nomads preferred to stay on the Southeastern border, while the majority became sedentary farmers ^53^. Three travel routes connect Ladakh with the Northern regions, and trade links have existed between Khotan, Yarkand, and Ladakh for thousands of years. Structures, rock-cut images, scattered sculptures, and carvings of Buddhist deities found at sites such as Mlbek, lc, Drss, ey, Saku, Padum, Tumel, Krtse, and Sni strongly reflect the Kashmiri tradition and are features of the second phase of the spread of Buddhism in Ladakh. Kashmiri has made contributions to the script of Ladakh, including the Boti script in Ladakhi, which is derived from the Kashmiri language ^54^.

Further archaeological and anthropological evidence of migration and trade movements includes the presence of wall fortifications. This provides insights into Ladakh’s past, cultural exchanges with Tibet and Central Asia, and involvement in Himalayan wars. The petrographic records of Ladakh describe the native flora and economy. The depiction of human masks from the 2nd and 3rd millennium BC that shows striking similarities to those from Southern Siberia and Northern Pakistan. Similarly, “tip-toe” creatures in later artworks suggest a connection to the Scythian societies of Central Asia from the 1st millennium BCE ^55^. Both examples suggest possible cultural mergers and trade movements in the Ladakh region. The genetic studies carried out so far on migration in the Trans-Himalayan region sometimes suggest contradictory hypotheses. On the one hand, the Himalayan border is considered a natural barrier for movement of people; on the other hand, it is considered a favorable corridor for human migration ^56^, but there is no solid genetic evidence to support such claims ^29^. Studies based on uniparental markers suggest high chromosomal diversity from East Asia (Uyghur, Han Chinese, and Nepalese), West and South Asia in terms of patrilineal markers, while the matrilineal studies suggest local ancestry and low diversity.

The Y chromosome haplogroup diversity in the region shows the presence of rare haplogroups such as Q, J1, R1a, L, and H, supporting the rich cultural assimilation mediated by males. Furthermore, the presence of these haplogroups suggests an early wave of migration from Central Asia and Southern Siberia around 1525 kya and a series of more recent waves of migration from Southern India around 10 kya ^28^. These results suggest the patrilocal nature of Ladakhi populations ^28,29^. The uniparental mitochondrial markers in the region support low but significant diversity, mainly from the Tibetan Plateau region and Northern China during the middle Holocene. This hypothesis is supported by the abundant occurrence of haplogroups M9, A21, and I4b ^57^. Several factors have led to the rich mix of ethnic diversity in the region, including the emergence of regimes and kingdoms such as the Kushan Empire, frequent Mughal invasions in the 14th to 16th centuries, and the slave trade in Central Asia in the 18th century ^10,29,32,58^. Analysis of the haplogroup base of Himalayan and adjacent populations shows high levels of genetic heterozygosity and massive interpopulation exchange, as well as a strong presence of shared ancestry between these groups ^56^.

We observed very clear dichotomy in the distribution of ancestral components between two Ladakh groups, termed LDKHA (Changpa) and LDKLA (Brokpa and Minero) from the genetic analysis of 80 unrelated Ladakh individuals. The comprehensive results show that the Ladakhi population is genetically split into two distinct clusters in frequency-based methods (Figs. 1b and c). The first cluster from the Leh region (Changpa) formed an extensive genetic cluster along the East Asian and Southeast Asian axis of the PCA. The second cluster, with two Ladakh groups (Minero and Brokpa), was closer to Northwest Indian high ANI groups ^59^ like Gujjar and Kashmiri Muslims in the Indian mainline and particularly among North Indian, Northwest Indian, Indo-European, and Nepalese Brahmin groups. Only a few Brokpa and Minero individuals and Nepalese Brahmins lie between two Ladakh clusters (Figs. 1b and c). Changpa’s (LDKHA) genetic ancestry is similar to that of the Tibetan and Sherpa populations, which is evident from the sharing of components in model-based clustering of admixture (Supplementary Fig. S2). On the other hand, the Minero and Brokpa (LDKLA) groups only showed genetic similarities with the Nepalese Brahmin and Kashmiri Muslim populations. Both groups shared South Asian components (Supplementary Fig. S2), but neither group shared the major ancestral components of East Asia or Southeast Asia. This may be a consequence of drift occurring in the Changpa population (presence of a distinct pink color component in Admixture plot) (Supplementary Fig. S2). Some of the earlier studies also recorded strong drift in South Asian populations, such as Kalash and Coorg ^60,61^. A high proportion of West Eurasian proportions was observed in Brokpa and Minero individuals. These patterns are more obvious in haplotype-based clustering, where Changpa (LDKHA) closely share clades with major East and Southeast Asian clade, whereas Brokpa and Minero (LDKLA) share clade with Indian Indo-European linguistic groups (Supplementary Fig S1). The same applies to the outgroup F3 and D statistics, where Changpa shares most alleles with East and Southeast Asian groups, while Brokpa and Minero share more alleles with Northwest Indian populations (Supplementary Table S1 and Fig. 1d and e). LDK-LA group shared higher number of IBD segments with Kashmiri Muslims, which probably reflects the major demographic changes during 14^th^ to 16^th^ century (Mughal impact). After religious conversions, movement further towards Kashmir and settlement of some of the Ladakh groups probably resulted into genetic assimilation of their component into Kashmiri groups. On the other hand, LDK-HA (Changpa) majorly share IBD segments with East Asian populations, reflecting shared genetic lineage with these groups, and probably reflecting the major gene flow during post-glacial from East Asia into the region ^30,62–64^.

These genetic findings are on par with archaeological evidence of possible diverse and multicultural migrations and settlements in the Ladakh region. The ceramic types found in the Leh region are consistent with the Tibetan type inhabited by the Changpa group ^8^, with higher affinity to East and Southeast Asia and genetically closer to the Tibetans and Sherpa groups. On the other hand, the other Ladakh regions (Changthang and Zanskar) were rich in corded pottery patterns, reflecting the West Eurasian influence ^8^. The West Eurasian influence on the genetic relationship of Brokpa and Minero confirms this type of corded pottery and illustrates very well the mixing of culture and genetics of the region. The genetic influence of ancient migrations from East Asia and Tibet has previously been well documented by studies that included mitochondrial markers with the prevalence of the East Asian M9a and Tibetan A21 haplotypes ^30^. The further sharing of the genetic component with Northwest Indian groups such as Gujjar and Kashmiri Muslims fits quite well with the historical accounts of the Mughal invasion and forced conversion of the Ladakh population, some of whom may have migrated and settled further downstream as Kashmiri Muslims. The Changpa people were likely more protected due to their geographical isolation (higher elevation) and Buddhist practices (protection from ruling classes). This can also be attributed to the higher strength of the founder event in Brokpa and Minero compared to Changpa. The estimated dates of the founder event in both groups (around the 22 generation in Changpa and around the 11 generation in Brokpa and Minero) also correspond to the historically documented Mughal invasion phase in the region ^15^ (around the 16th century). Admixture model inference and demographic modelling suggest that Changpa or LDKHA are probably derived from Tibetan like sources with very recent admixture from French-like (Kashmiri Muslim) source. On the other hand, LDKLA (Brokpa and Minero) are lineages derived from an admixture event between Kashmiri Muslim (or French-like source) and Tibetan with additional admixture recently from Tibetan. In demographic modelling only slight decrease in effective population size of LDKLA was observed. Our simulation-based analysis clearly suggests that higher distribution of genome wide RoH segments in Ladakh-LA group may be consequence of such stronger founder event in them compared to Ladakh-HA group. Although, higher mean length of RoH at 1Kb window in Ladakh-LA may also be resultant of geographic isolation (higher altitude). Selection scan-based genetic signatures in the Changpa group showed known signatures of high-altitude adaptation, with some broadly categorized selection markers among immunity-related genes. Similar genetic signatures were not observed in Brokpa and Monpa, but they did exhibit immunity-related genetic signatures again pointing out their distinct genetic lineage.

## Data availability

Genotype data of 80 individuals from Ladakh is publicly available with free access from following Zenodo accession id: https://doi.org/10.5281/zenodo.11217308.

## Acknowledgements

We thank all the study participants, who volunteered in this study. NR thank the Director, Birbal Sahni Institute of Palaeosciences for scientific support and Department of Science and Technology for CRG grant (CRG/2021/006762) from SERB. KT was supported by J C Bose Fellowship (JCB/2019/000027) from the Science and Engineering Research Board (SERB), Department of Science and Technology, Government of India.

## Author contributions

Conceptualisation: K.T. and N.R.; sample collection and genotyping: N.P., R.R., S.S., S.R.; data analysis and visualisation: L.K.; K.T., N.R., S.S. significantly contributed to results interpretation; L.K. wrote the original draft, with contributions from R.R., B.A., S.K., A.D. and S.K.; K.T., N.R., S.S., and S.R. revised the manuscript and contributed to the final version.

## Ethics declarations

### Conflict of interest

None.

### Ethics approval

The study was approved by Institute Ethical Committee of BSIP, Lucknow, India.

### Informed consent

Informed written consent was obtained from all the participants involved in the study.

## Notes

### Competing Interest Statement

The authors have declared no competing interest.

### Summary of Updates

This manuscript version contains some additional analysis based on the newly added demographic modeling of the Ladakh populations.

