## Supplementary text for "Genetic architecture and adaptation of Ladakh highlanders of trans-Himalayas"

**Supplementary Fig 1.**

**A.** Best fitted demographic model inferred from Moments and corresponding model parameters for LDKHA group

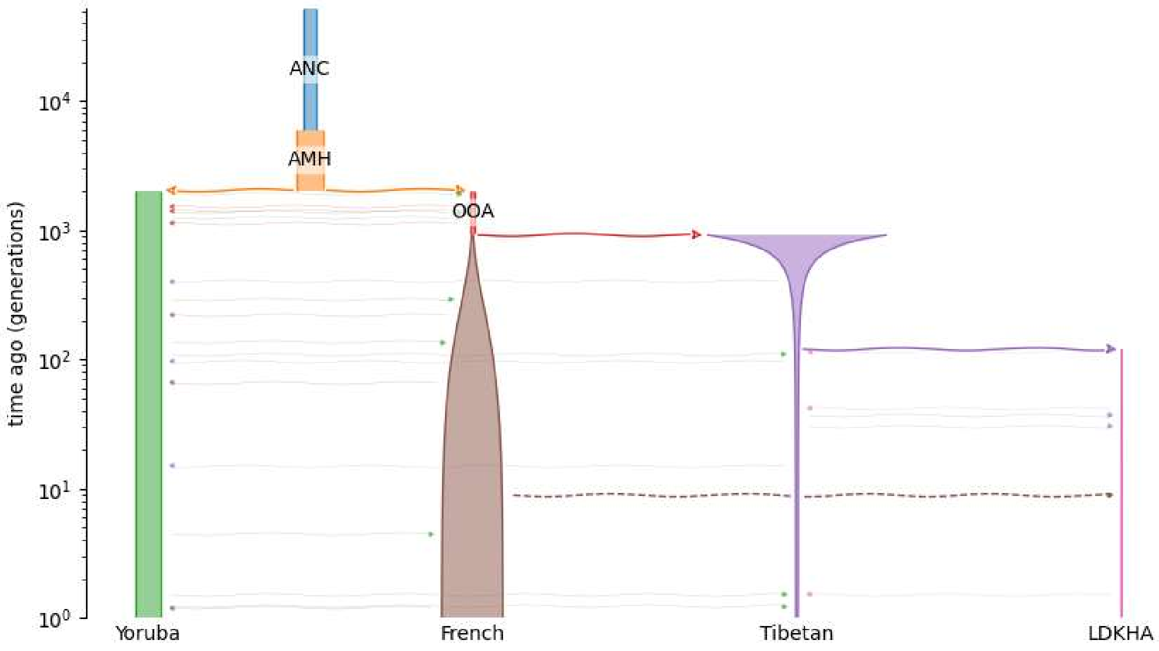


| Log-likelihood: -114400.79498136348 | | | | |
|---|---|---|---|---|
| **Parameter** | **Description** | **value** | **2.5%** | **97.5%** |
| Ne | Tibetan initial size | 9.99e+04 | 79421 | 120423 |
| NeF | Tibetan final size | 1.36e+03 | 1212.4 | 1512.81 |
| NA | LDKHA initial size | 1e+02 | nan | nan |
| NF | LDKHA final size | 4.62e+02 | 363.411 | 559.652 |
| M_Tib_LDKHA | migration rate b/w Tibetan and LDKHA | 0.0379 | 0.0357717 | 0.0400702 |
| Pulse1 | pulse time from French to LDKHA | 8.89 | 7.57891 | 10.1925 |

**B.** Multinomial comparison between 4d model and data and Anscombe residuals

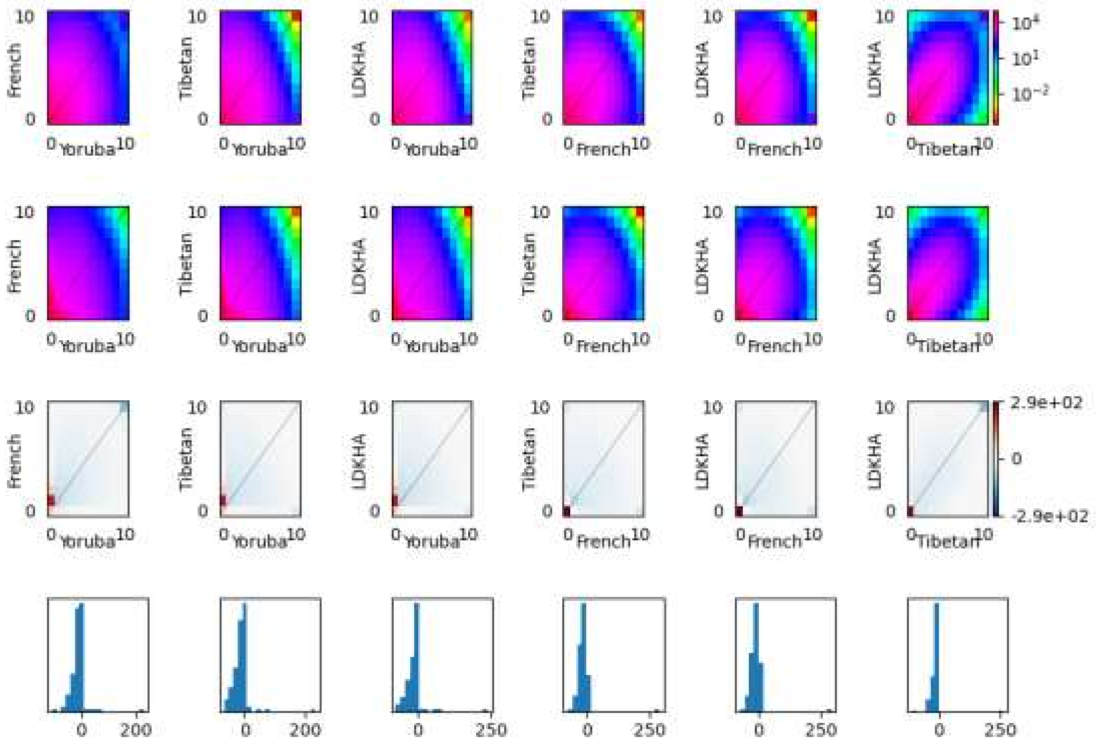

**Supplementary Fig 2.**

**A.** Best fitted demographic model inferred from Moments and corresponding model parameters for LDKLA group

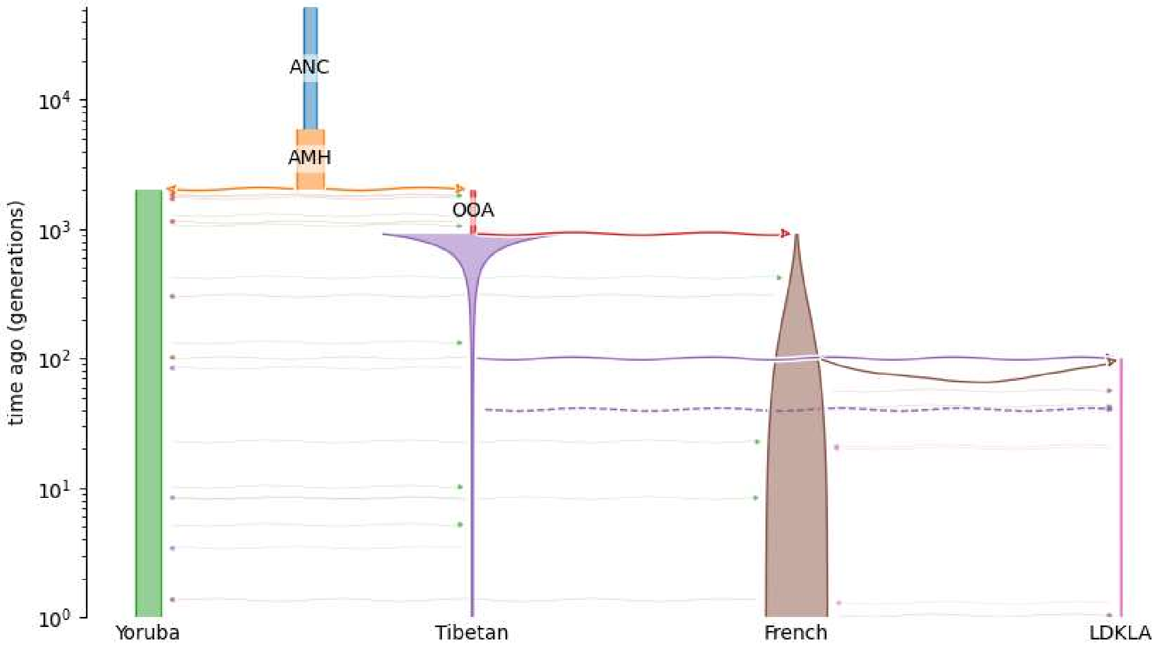


| | | | | |
|---|---|---|---|---|
| Log-likelihood: -121125.18700491334 | | | | |
| Parameter | Description | value | 2.5% | 97.5% |
| Ne | Tibetan initial size | 9.96e+04 | 92454.8 | 106720 |
| NeF | Tibetan final size | 7.6e+02 | 742.521 | 776.526 |
| NA | LDKLA initial size | 6.98e+02 | nan | nan |
| NF | LDKLA final size | 4.39e+02 | nan | nan |
| M_French_LDKLA | migration rate b/w French and LDKLA | 0.00255 | 0.00244887 | 0.00264396 |
| Pulse1 | pulse time from Tibetan to LDKLA | 40 | 36.0322 | 44.8412 |

**B.** Multinomial comparison between 4d model and data and Anscombe residuals

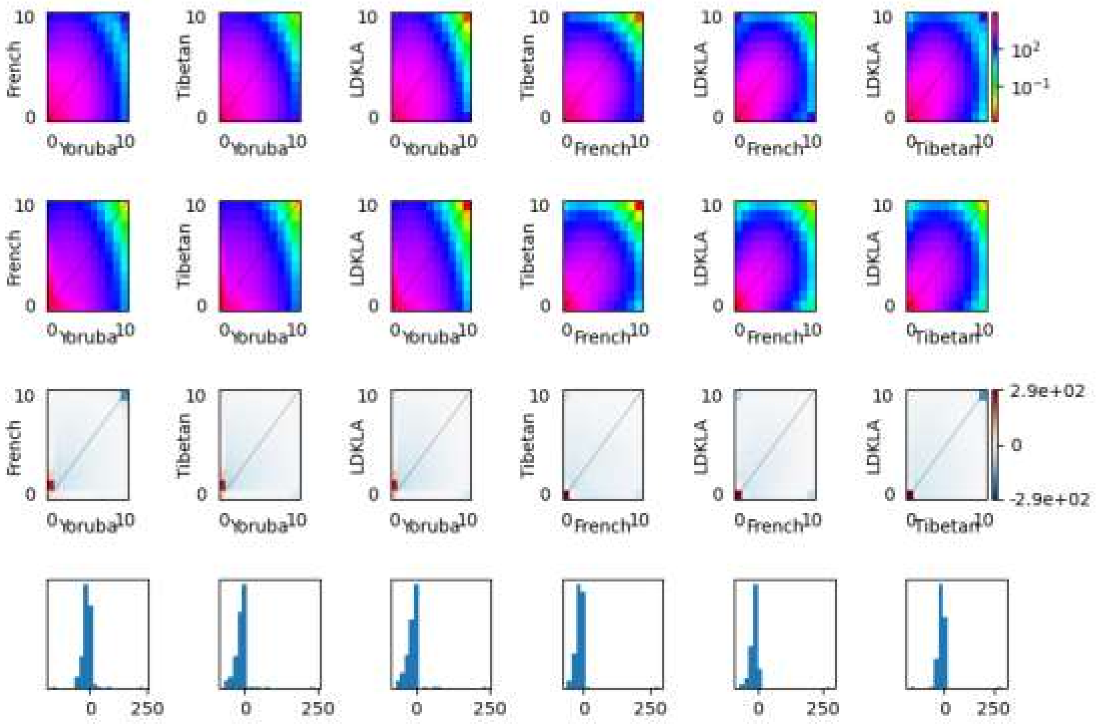

**Supplementary Fig 3.** Genetic simulations to test the effect of founder events typical of both the Ladakh clusters on the genome-wide RoH distribution. The genetic simulation scenarios to replicate the Ladakh-LA **A.** and Ladakh-HA **B.** founder events. Estimated strength and timing of founder event from simulated data from Ladakh-LA **C.** and Ladakh-HA **D.** scenarios. RoH distribution from simulated data at window of 1kb **E. and** RoH distribution at window of 5kb **F.**

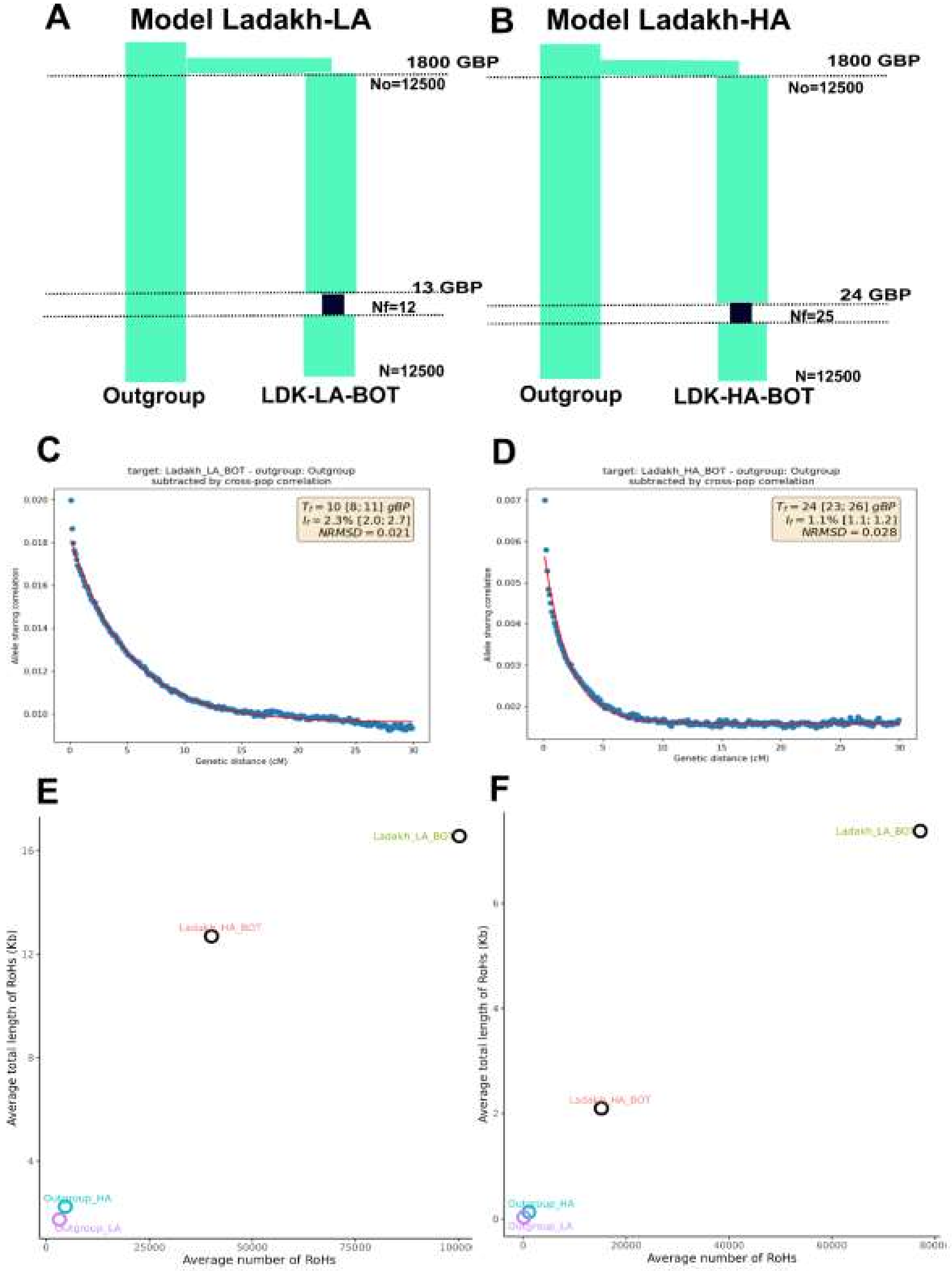

**Supplementary Fig 4**. Heatmap of Chunk count sharing matrix of ChromoPainter run with modern Eurasians

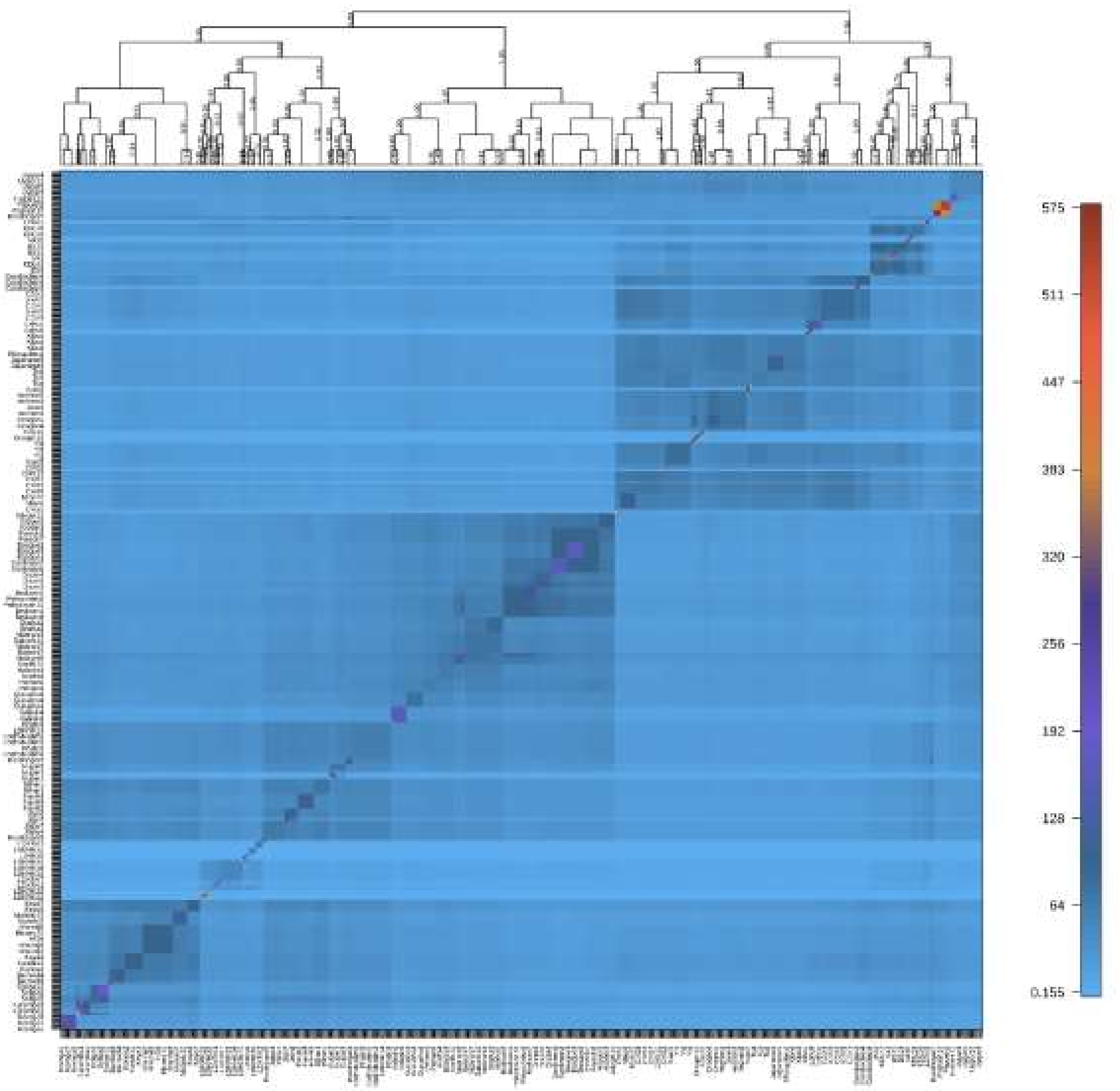

**Supplementary Fig 5.** Genome-wide RoH distribution at 1kb window **A.** and 5kb window size **B.**

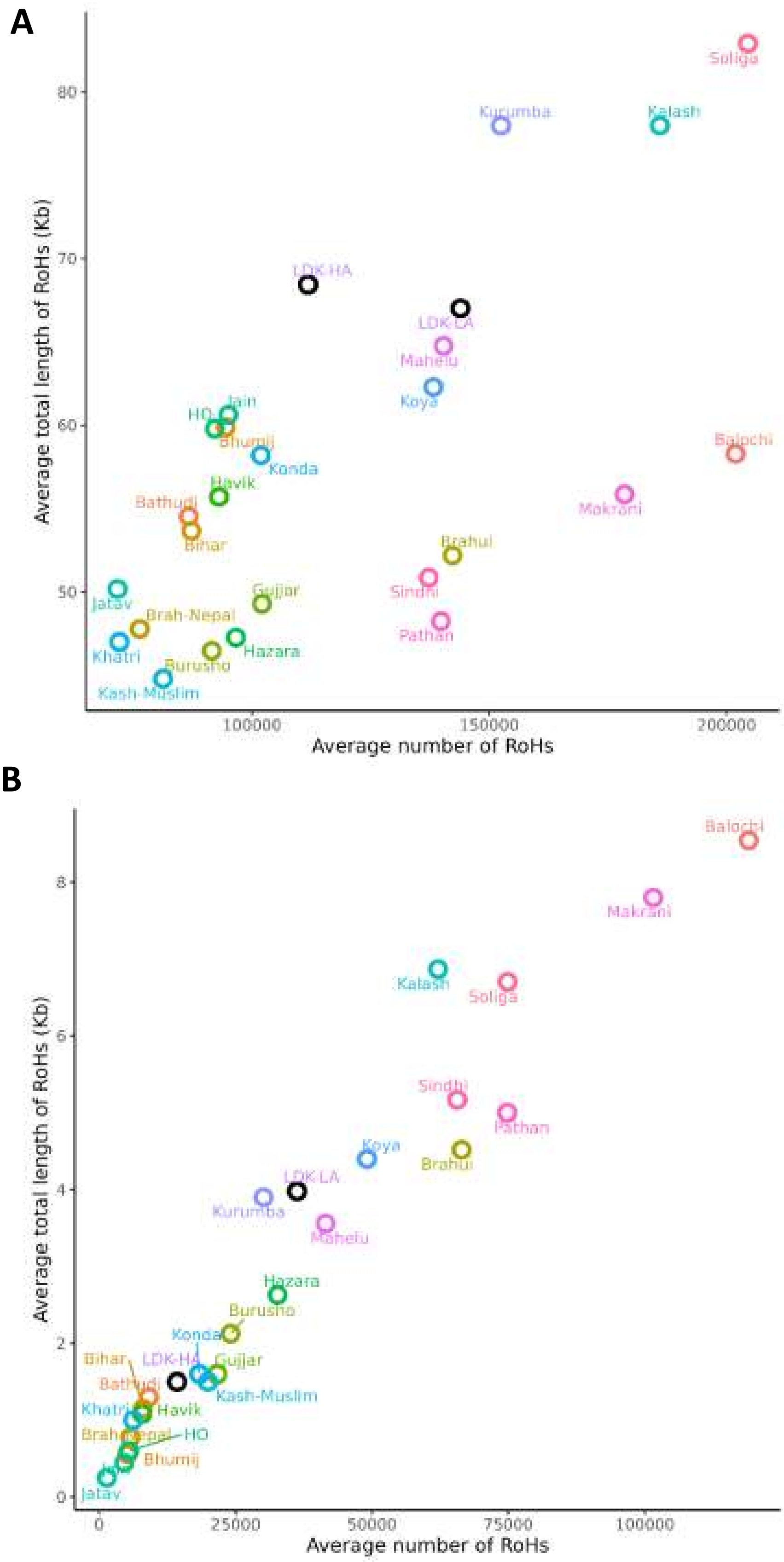

**Supplementary Fig 6. A.** ML tree constructed by TreeMix, **B.** Residual plot

**A**

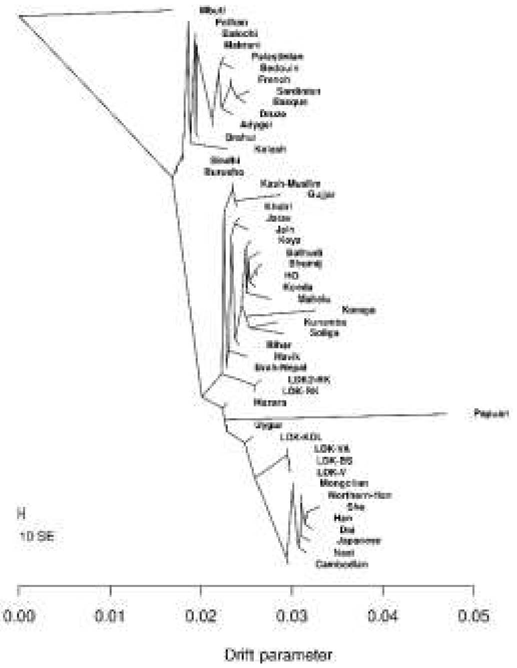


**B**

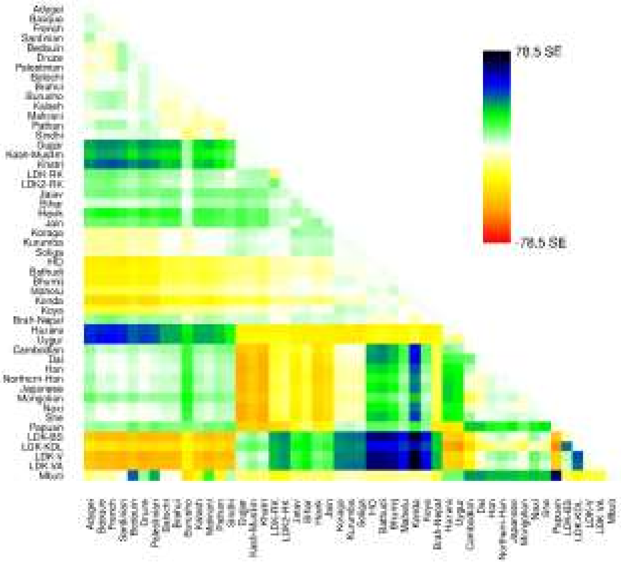

**Supplementary Fig 7**. Manhattan plot showing loci under selection in population **A.** Ladakh-HA and **B.** Ladakh-LA.

**A**

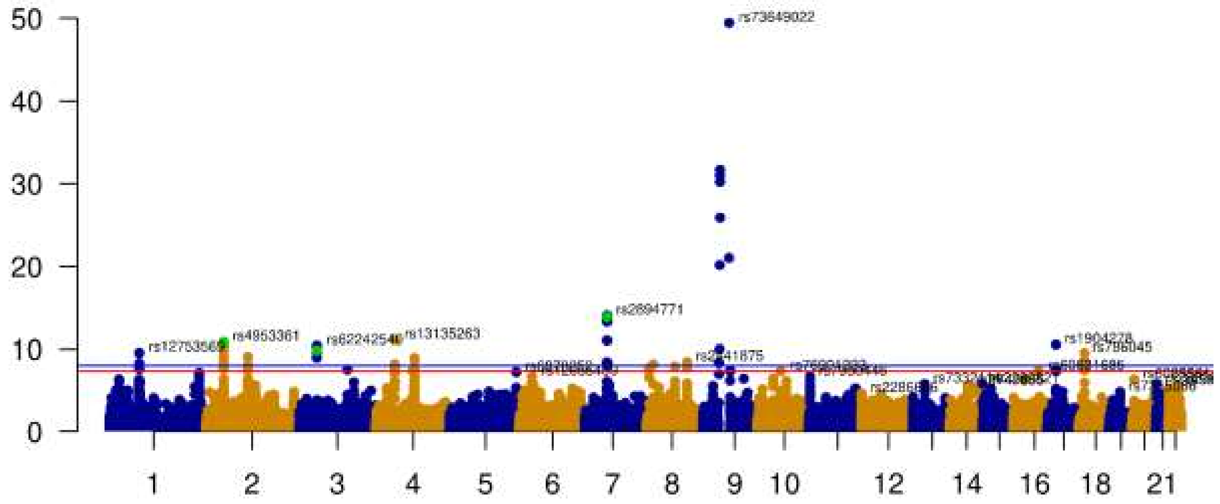


**B**

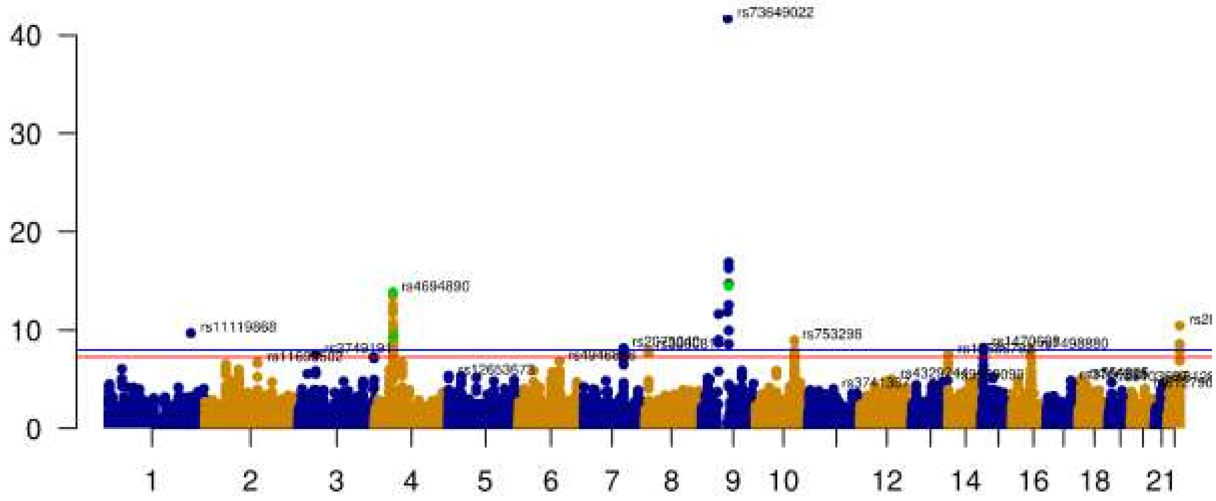
